# Key Factors in the Cortical Response to Transcranial Electrical Stimulations—A Multi-Scale Modeling Study

**DOI:** 10.1101/2021.10.06.463305

**Authors:** Hyeyeon Chung, Cheolki Im, Hyeon Seo, Sung Chan Jun

## Abstract

Transcranial electrode stimulation (tES), one of the techniques used to apply non-invasive brain stimulation (NIBS), modulates cortical activities by delivering weak electric currents through scalp-attached electrodes. This emerging technique has gained increasing attention recently; however, the results of tES vary greatly depending upon subjects and the stimulation paradigm, and its cellular mechanism remains uncertain. In particular, there is a controversy over the factors that determine the cortical response to tES. Some studies have reported that the electric field’s (EF) orientation is the determining factor, while others have demonstrated that the EF magnitude itself is the crucial factor. In this work, we conducted an in-depth investigation of cortical activity in two types of electrode montages used widely—the conventional (C)-tES and high-definition (HD)-tES—as well as two stimulation waveforms—direct current (DC) and alternating current (AC). To do so, we constructed a multi-scale model by coupling an anatomically realistic human head model and morphologically realistic multi-compartmental models of three types of cortical neurons (layer 2/3 pyramidal neuron, layer 4 basket cell, layer 5 pyramidal neuron). Then, we quantified the neuronal response to the C-/HD-tDCS/tACS and explored the relation between the electric field (EF) and the radial field’s (RF: radial component of EF) magnitude and the cortical neurons’ threshold. The EF tES induced depended upon the electrode montage, and the neuronal responses were correlated with the EF rather than the RF’s magnitude. The electrode montages and stimulation waveforms caused a small difference in threshold, but the higher correlation between the EF’s magnitude and the threshold was consistent. Further, we observed that the neurons’ morphological features affected the degree of the correlation highly. Thus, the EF magnitude was a key factor in the responses of neurons with arborized axons. Our results demonstrate that the crucial factor in neuronal excitability depends upon the neuron models’ morphological and biophysical properties. Hence, to predict the cellular targets of NIBS precisely, it is necessary to adopt more advanced neuron models that mimic realistic morphological and biophysical features of actual human cells.

## 1. Introduction

Noninvasive brain stimulation (NIBS) has garnered increasing attention from many researchers as a potential treatment for neurologic and psychiatric disorders and a neuromodulation technique in neuroscience brain research [1]. Briefly, NIBS is categorized into transcranial electrode stimulation (tES) and transcranial magnetic stimulation (TMS). tES is a technique that modulates cortical activities by delivering weak electric currents through scalp-attached electrodes. There are several tES techniques—transcranial direct current stimulation (tDCS), transcranial alternating current stimulation (tACS), transcranial random noise stimulation (tRNS), electroconvulsive therapy (ECT), and so on [2]. Among these, tDCS/tACS are of particular interest to us; they modulate cortical activity by passing an electrical current through the skull from electrodes attached to the scalp [3]. While ECT invokes neuronal activation directly with a strong intensity current, tDCS and tACS modulate spontaneous neural activity at low intensity (0.5-2mA). The electric field (EF) tDCS induces by delivering direct electrical current is subthreshold, and therefore, tDCS may modify neuronal transmembrane potentials and thus influence the neurons closer to their EF threshold without depolarization [4]. In contrast, tACS delivers an alternating electrical current and causes the neuronal transmembrane to fluctuate according to the currents; it is reported that applying a lower frequency of alternating current causes greater polarization than does a higher frequency [3].

NIBS has been applied widely in neuromodulation and treatments for neurologic and psychologic disorders according to their specific characteristics. However, despite the growing cases of applications and increased attention to NIBS, the brain regions affected and neuronal responses’ cellular mechanism remain undetermined clearly. Thus, experimental and computational studies have been conducted to attempt to resolve this uncertainty at the cellular (microscopic) and cortical (macroscopic) levels. At the cellular level, both simulations and experimental studies have found that the intrinsic neuronal morphology is a crucial factor in the neuronal response to an electrical stimulus (with respect to the threshold, membrane polarization, and firing patterns) [4–15]. In particular, *in vitro* studies have revealed that neuronal morphology affects such firing properties as threshold, sensitivities to EF, and firing patterns [6–9, 11, 15]. Different morphologies cause variations in neuronal responses to stimulation, even within a cell type [7, 15]. Further, morphological parameters, such as diameter, length, and the degree of arborization’s effects on neuronal responses have been investigated through simulations that modified morphological parameters [5], and a simulation study found that dendritic structures affected not only firing properties, but also action potential efficiency [9]. Thus, it is essential to incorporate realistic neuron models to understand the mechanism of actual neuronal responses to NIBS more deeply.

To achieve a better estimation of neurons’ response to the stimulation, it is necessary to map morphologically realistic neuronal models to a head model that reflects realistic anatomy and consider the EF’s spatial intensity (macroscopic level) and the neurons’ morphological/biophysical properties (microscopic level). This is because cortical geometry is of utmost importance to understand inter-subject variability because of its effect on the EF’s spatial distribution [16–24], while the EF applied to a neuron depends upon the targeted neuron’s location in a head model and the anatomical head geometry. Thus, to estimate neuronal responses better, it is important to consider the neurons’ biophysical and morphological properties and their spatial location within the brain. However, the *in vivo* recording of EF in several positions during stimulation is difficult, as it requires invasive recording electrodes, i.e., implanting electrodes on the cortex [4, 25–27]. As an alternative, a simulation study plays a key role in estimating the spatial intensity of activation and the neurons’ distinguishable responses to the stimulation. Recent attempts have been made to consider both levels by constructing a multi-scale model that combines realistic multi-compartmental neuron models and an anatomically realistic head model [28–30]. However, despite these efforts, uncertainty yet remains because of the technical challenges in incorporating both neuron/head models and a limited understanding of neuromodulation’s cellular mechanism.

Moreover, there is a controversy surrounding the determining factors in the neuronal response to NIBS, although various attempts have been made to determine the key factors in neuronal responses to such NIBS techniques as TMS and tDCS. Some studies have demonstrated that EF orientation is a key factor [16, 21, 28, 31–36], while others have argued that EF magnitude is a crucial factor [19, 29, 37, 38]. In particular, a TMS study suggested that the EF’s strength affects the neurons’ excitation threshold highly [19], but other studies have observed that cortical activation is sensitive to orientation [34, 35]. Further, in the case of tDCS studies, two divergent results have been reported [16, 32, 33, 36, 37]. In the absence of consensus, various multi-scale models for TMS have been introduced. Particularly, a recent multi-scale model that used realistic neurons with arborized axons reported that the magnitude of the TMS-induced EF affected cortical excitability [29]. However, most simulation studies of NIBS are constrained to TMS, and there has been little investigation of the cellular mechanisms in tES. Although our group conducted multi-scale modeling research of tES [28], we used a neuron model with an idealized straight axon. However, the importance of incorporating realistic axons in a simulation study for TMS have been reported [14, 29, 31]. Thus, investigating the cellular mechanism for tES with realistic and arborized axons is necessary to eliminate the remaining uncertainty and resolve the question—What is the determining factor in the neuronal response to NIBS?

The absence of studies of tES and unresolved controversy motivated us to conduct an in-depth investigation of the realistic cortical excitability that tES causes (with a focus on tDCS and tACS). To do so, we constructed a multi-scale model for conventional (C-) and high-definition (HD-) tES by coupling an anatomically realistic head model with multi-compartment neuron models that retain a realistically detailed morphology from the dendrites to the axon collaterals. These multi-scale models reflected the cortical neurons’ morphological/biophysical properties and the brain model’s cortical geometry. This reflection of both the microscopic and macroscopic levels allowed us to quantify the neuronal response to a realistic tES-induced EF calculated in the anatomically realistic head model by measuring the cortical neurons’ excitation threshold. We observed neuron’s firing properties to a strong stimulation for an in-depth investigation of neurons, because the neuronal response to weak stimulation (<1-2mA) is linear polarization. Moreover, we investigated neuronal morphology and its effect on neurons’ response to tES by rotating neurons within the cortex. As 10Hz-tACS is used to enhance motor learning [39, 40] and motor memory [41] because of its capability to target a specific oscillation, we investigated 10Hz tACS as well as tDCS. Thereafter, we investigated the extent to which the excitation threshold is correlated with EF and RF (radial component of EF) to resolve the controversy over neuronal responses’ determining factors. Our investigation was the first attempt to incorporate realistic axons in an electrical stimulation study. As a result, we identified the brain region C-and HD-tES affected, the cortical neurons’ excitation thresholds, the importance of neurons’ morphology, compared the results of tDCS and tACS, the degree of correlation between EF and RF magnitude, and the excitation threshold.

## 2. Methods

To create a multi-scale model for C-/HD-tES, we mapped realistic neuron models onto a realistic head model. However, it requires enormous computational resources to simulate all neuron models over the entire brain. Thus, we considered a region of interest (ROI) and confined the neuron models within the ROI of the head model. In this work, our chosen ROI was the hand knob area that consists of the precentral gyrus and some part of the postcentral gyrus. We calculated the realistic EF distribution under C-/HD-tES montages (Figures 1a and 1b) and applied an EF to neurons within the ROI. Then, we quantified the neuronal response to the stimulation by examining the excitation threshold, which is defined as the minimum stimulus amplitude that evokes an action potential in the soma.

**Figure 1.**
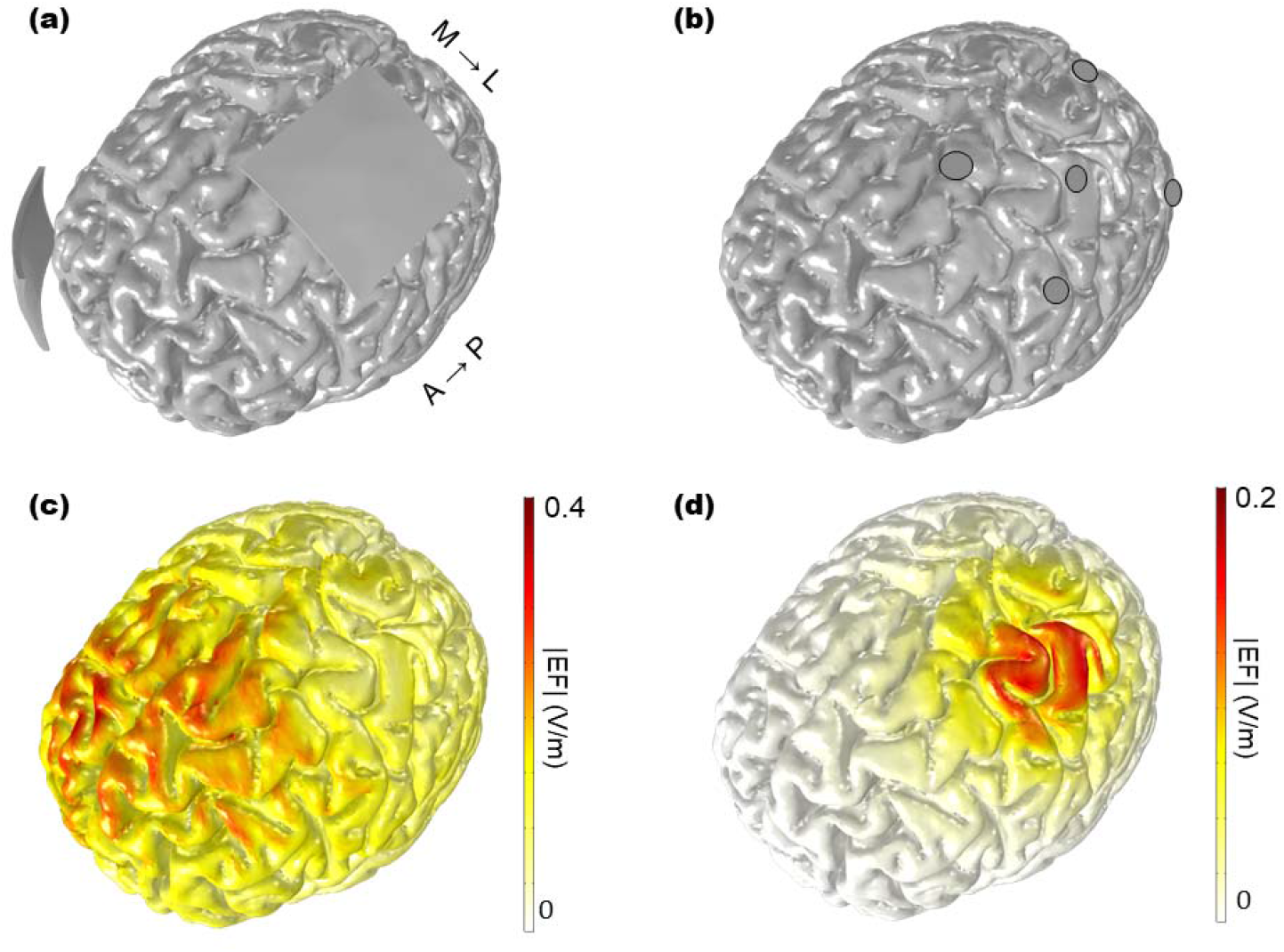
tES montage and distributed EF. C-tDCS/tACS (C-tES) (a) and HD-tDCS/tACS (tES) (b) electrodes were placed over the ROI. The EF distributions (in magnitude) of C-tES (c) and HD-tES (d) are illustrated on the brain.

### 2.1. Realistic head model and EF calculation

We introduced an anatomically realistic volume conduction head model provided by SimNIBS v. 2.0 that reflects anatomical geometry fully [42]. This head model has approximately 12.2 million elements and approximately 2 million nodes, and consists of five layers: gray matter; white matter; cerebrospinal fluid (CSF); scalp, and skull. We considered two types of montages, C- and HD-tES. For C-tES, two patch-type electrodes 5mm × 5mm and 2mm thick were attached to the scalp: One active (anode) electrode was attached over the ROI and the other over the frontal area as the reference (cathode) (Figure 1a). With respect to HD-tES, five disc-type electrodes (4mm radius, 2mm thick) were attached to the scalp with 2mm thick CCNY-4 gels according to the 4 × 1 electrode configuration (Figure 1b), including one active (anode) electrode and four reference (cathode) electrodes surrounding the active electrode [23]. Each tissue layer of the head model was assigned isotropic electric conductivity values (in S/m) [23, 43, 44], scalp: 0.465; skull: 0.01; CSF: 1.654; gray matter: 0.276, white matter: 0.126; patch-type electrode: 1.40; disc-type electrode: 5.8 × 10^7^, and gel: 0.30. More modeling details can be found in [28].

After constructing the volume meshes for the head model, the Laplace equation ∇· (σ∇V) = 0 (V: potential, σ: electrical conductivity) was solved by FEM implemented in COMSOL Multphysics (v. 5.2a, COMSOL, Inc., Burlington, MA, USA) to compute the EF distributions under the C- and HD-tES montages. The conjugate gradient method was used with a relative tolerance of 1 × 10^−6^ with a precondition of an algebraic multigrid. In this calculation, the following boundary conditions were used: Normal current density (inward current flow) was applied to the external surface of the active electrode (V = constant); ground was applied to the exposed surface of the reference electrode (V = 0), and the remaining exposed surfaces were configured as an electric insulator [23, 28]. Finally, we set a constant current stimulation of 1mA passing through the active electrode and calculated the EF distribution over the entire head model. In addition, we measured the RF, which is a directional component of EF that is normal to the cortical surface. The RF was calculated by 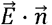, in which 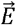 is a calculated EF, and 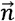 the inward normal vector of the triangular element on the cortical surface.

### 2.2 Realistic Neuron Models

We adopted multi-compartmental models of a layer 2/3 pyramidal neuron (L2/3 PN), layer 4 large basket cell (L4 LBC), and layer 5 pyramidal neuron (L5 PN), which are morphologically and physiologically realistic (Figure 2), and are available from ModelDB (Model 241165) [45]. The models contain detailed and realistic morphologies from dendrites to arborized axons and their biophysical properties have been validated experimentally [7, 46]. The neuron models are a modified version of the Blue Brain Project’s multi-compartmental models [46, 47] implemented in the NEURON simulator environment [48]. The modifications involved myelinating the axonal arbors, scaling ion channel kinetics, and assigning ion channel characteristics to all of the axons’ arborized points [7], as well as scaling morphologies to account for age and species variations [7, 29]. In particular, with respect to L5 PN, axon collaterals were extended to compensate for truncation of axons in the Blue Brain Project’s slicing process [46, 47]. For more details of modifications, refer to [7, 29].

**Figure 2.**
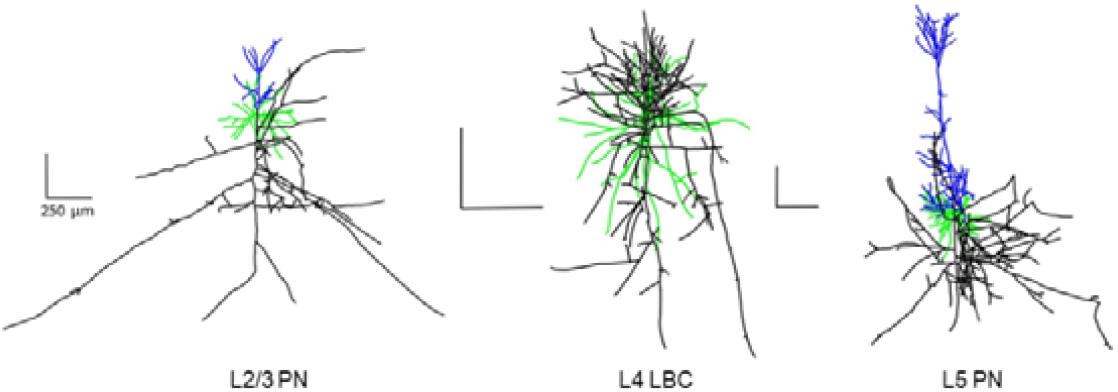
Multi-compartment neuron models: layer 2/3 pyramidal neuron, layer 4 large basket cell, and layer 5 pyramidal neuron. Morphologies are colored to indicate axon (black), apical dendrite (blue), and basal dendrite (green).

We chose one neuron from each of the three cell types (L2/3 PN, L4 LBC, L5 PN) in an original set of neuron models that consist of five clones of five cell types. Representation of these different cell types is important to account for pyramidal neurons and basket cells’ various crucial functions in transmitting information across cortical layers. While pyramidal neurons process and deliver information, interneurons regulate the information processing at the local level [49]. Basket cells operate as an intercolumnar inhibitor within their cortical layers, and are one type of interneurons that are categorized into various cell types according to function and morphology [49]. The neuron model consisted of a series of sections divided into compartments, and these multi-compartmental neurons have been modeled by the cable equation [50]. Thus, ionic currents within various neuron models are represented spatially as their values at the center points of the compartment. Further, the axial current within the neuron is calculated by the voltage difference between each compartment’s center point.

### 2.3 Multi-scale modeling

To construct a multi-scale model for C-/HD-tES, we incorporated the realistic head model and the three types of neuron models, and populated neurons in a specific layer of the cortex in the ROI chosen (Figure 3a-3b). Each neuron was aligned according to each face of the tetrahedral meshes and at a particular layer depth, with the dendrite-to-soma axis perpendicular to each center point of the triangular face. During the alignment, we hypothesized two cases: neurons’ non-rotation and rotation. In the latter case, we rotated randomly around the dendritic-to-soma axis (Figure 3b-3c). The neurons’ rotation angles were chosen randomly among 0, 60, 120, 180, 240, and 300^0^ in accordance with Aberra’s report [29]. To achieve a better simulation, we created three sets with different random rotation trials, calculated their thresholds, and obtained the mean thresholds. Thus, we note that the excitation threshold for the rotated case is the mean threshold for the three cases with different random rotation trials. We delineated between the cortical layers and soma of the L4 LBC, L2/3 PN, and L5 PN, which were placed in distinct layers according to their median depth, which reflected *in vivo* experimental data [51]. Finally, a total of 3,132 neurons of each cell type was populated in the ROI.

**Figure 3.**
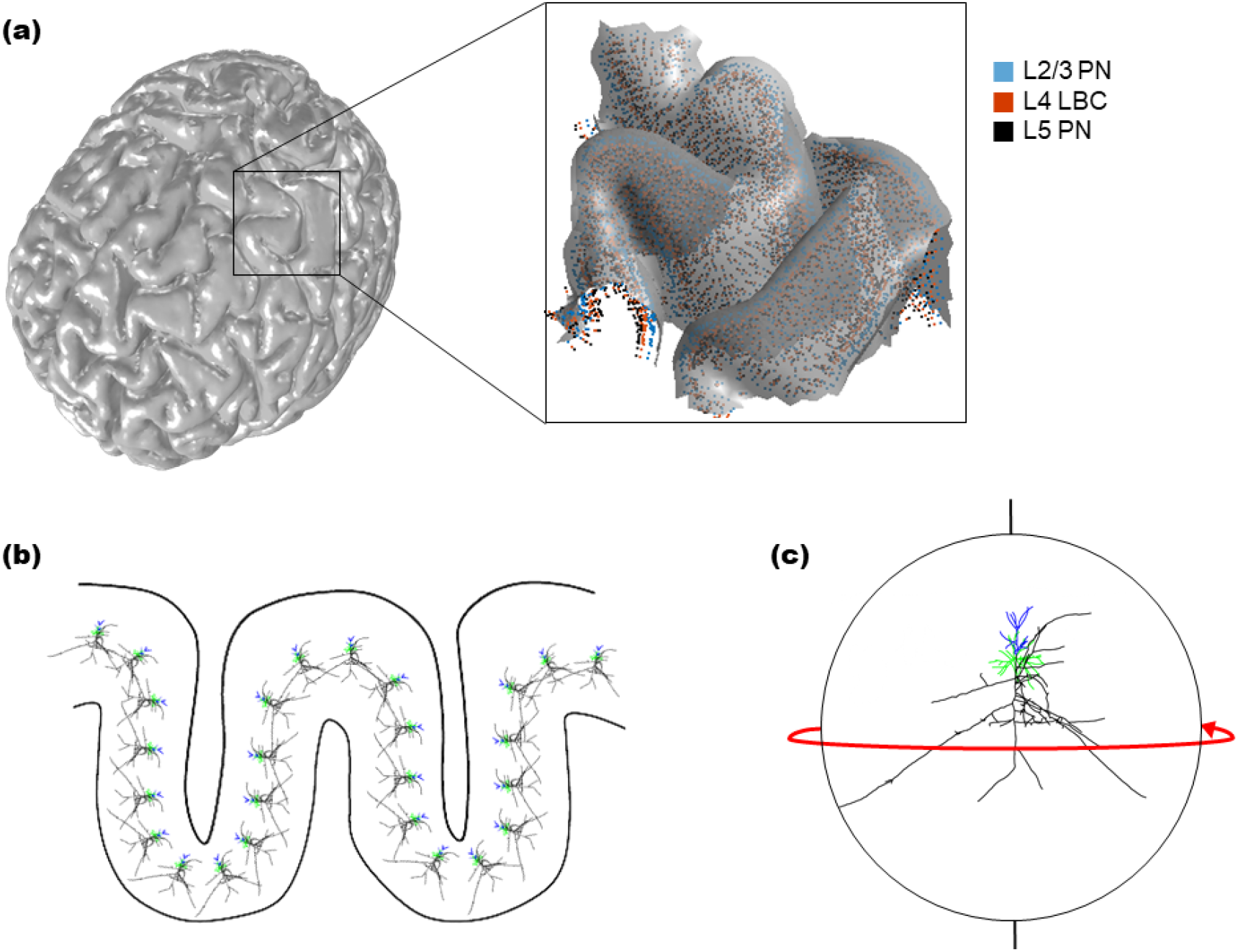
Schematic view of neurons populated in the ROI. (a) The hand knob area (ROI) on the precentral gyrus and its adjacent part of the postcentral gyrus (left small black box). Each neuron’s soma populated in the ROI is depicted as a dot and an enlarged view of the ROI is illustrated in the square inset. (b) Schematic view showing the way the L2/3 PNs are placed in the gray matter of the gyrus. In our model, neurons are relatively smaller and denser than those shown in this figure. (c) In the rotated case, neuron models are rotated along the dendrite-to-soma axis (line from top to bottom in the figure) and the rotation angle was one among 0, 60, 120, 180, 240, and 300°.

To simulate a neuron’s response to electrical stimulation within a head model, we applied a realistic tES-induced EF to the neuron models. The spatially variable potential field was calculated at each center point of the neuron models’ compartments located in the ROI based upon the simulated EF distribution in the head model. We used an extracellular method in the NEURON simulation environment (v. 7.7) [48] to map each potential value onto the center point of each neuron compartment, and simulated the neuronal response to the electrical stimulation of a 100ms monophasic square waveform (tDCS) and a 10Hz sinusoidal waveform (tACS). We observed that the threshold converged after 20-30ms when stimulation of a duration ranging from 1 to 100ms was simulated, which is consistent with a study reported previously [52], and thereby, we stimulated neurons with tDCS and tACS for 100ms. After mapping and computing the potential field, an excitation threshold for each populated neuron was determined to quantify the neuronal response. The excitation threshold, the minimum stimulus required to induce an action potential, was measured in the soma by scaling the potential field magnitude calculated in the head model with a stimulus amplitude from 1 to 1000mA. Activation of a neuron was defined as the somatic membrane potential crossing 0mV (measured by the APCount of point process in the NEURON simulation environment [48]).

## 3. Results

### 3.1. Uniform field

To gain a better understanding of the neuronal response to the EF, we calculated a uniform EF and applied it to the neurons. To calculate the EF, we used COMSOL Multiphysics (v. 5.2; Burlington, MA) to construct a three-dimensional cube model that covered all neuronal models. Each neuron’s soma was mapped onto the center of the cube model. Then, we observed the neurons’ threshold under uniform EF and the membrane polarization of the apical dendrites, soma, and axon terminals in L2/3 PN, L4 LBC, L5 PN. The threshold for 100ms radial field (RF) stimulation (current flow was applied parallel to the dendrite-to-axon axis) was 6 (L2/3 PN), 19 (L4 LBC), and 12mV/mm (L5 PN), respectively. The threshold for 100ms tangential field (TF) stimulation (current flow was applied perpendicular to the dendrite-to-axon axis) was 12 (L2/3 PN), 21 (L4 LBC), and 18mV/mm (L5 PN), respectively. From these thresholds, we inferred that the EF activated our neuron model both radially and tangentially, and the thresholds differed among cell types. The thresholds of L4 for RF and TF were higher than for other cell types, and the threshold difference between RF and TF in the L2/3 PN was larger.

### 3.2. EF and threshold distribution

We delivered a 1mA stimulus of direct current (tDCS) and alternating current (tACS) to the targeted ROI for 100ms via scalp-attached active electrodes for C- and HD-tES. With respect to the EF distribution, HD-tES yielded more focality, but less strength, while C-tES was stronger, but diffused (Figure 1c-1d). This pattern is consistent with those in previous studies [23, 28, 53–56]. C-tES’s maximum EF in the entire brain was 0.67 V/m, and HD-tES’s was 0.21 V/m. Within the ROI, C-tES’s maximum EF was 0.25 V/m and HD-tES’s was 0.21 V/m, while C-tES/HD-tES’s minimum EFs were 0.05 and 0.02 V/m, respectively. With respect to the distribution, for both C- and HD-tES, a higher EF magnitude was found in the crown (the top of the gyrus) and a lower magnitude was observed in the bank (a deeper region of the sulcus in both the pre-and post-central gyrus of ROI).

Interestingly, a similar pattern of EF distribution was found in the three neuron models’ threshold distributions. First, with respect to tDCS stimulation, the thresholds of HD-tDCS were higher than those of C-tDCS, as C-tDCS generated a higher EF than HD-tDCS in both the unrotated and rotated cases (Figures 4a and 4c). In the unrotated case, the median thresholds of C-tDCS were 33, 133.5, and 117 mV/mm for L2/3 PN, L4 LBC, and L5 PN, respectively. Further, the median thresholds of HD-tDCS in the unrotated case were 49, 181, and 152 mV/mm for L2/3 PN, L4 LBC, and L5 PN, respectively. However, in the rotated case, the median thresholds for C-tDCS were 34.7, 141.3, and 125 mV/mm for L2/3 PN, L4 LBC, and L5 PN, while those for HD-tDCS were 54.3, 185.5, and 165.7 mV/mm for L2/3 PN, L4 LBC, and L5 PN. Second, with respect to tACS stimulation, the thresholds of C-/HD-tACS were higher than C-/HD-tDCS overall, and the thresholds of HD-tACS were higher than those of C-tACS (Figures 4b and 4d). In the unrotated case, the median thresholds of C-tACS were 55, 165, and 127 mV/mm for L2/3 PN, L4 LBC, and L5 PN, respectively. In addition, the median thresholds of HD-tACS in the unrotated case were 76, 214, and 159 mV/mm for L2/3 PN, L4 LBC, and L5 PN, respectively. However, in the rotated case, the median thresholds for C-tACS were 55.5, 177.7, and 132 mV/mm for L2/3 PN, L4 LBC, and L5 PN, and those for HD-tACS were 76, 214, and 169.7 V/mm for L2/3 PN, L4 LBC, and L5 PN, respectively. Overall, the stimulation waveform’s (tDCS/tACS) threshold varied somewhat, but rotation caused small changes.

**Figure 4.**
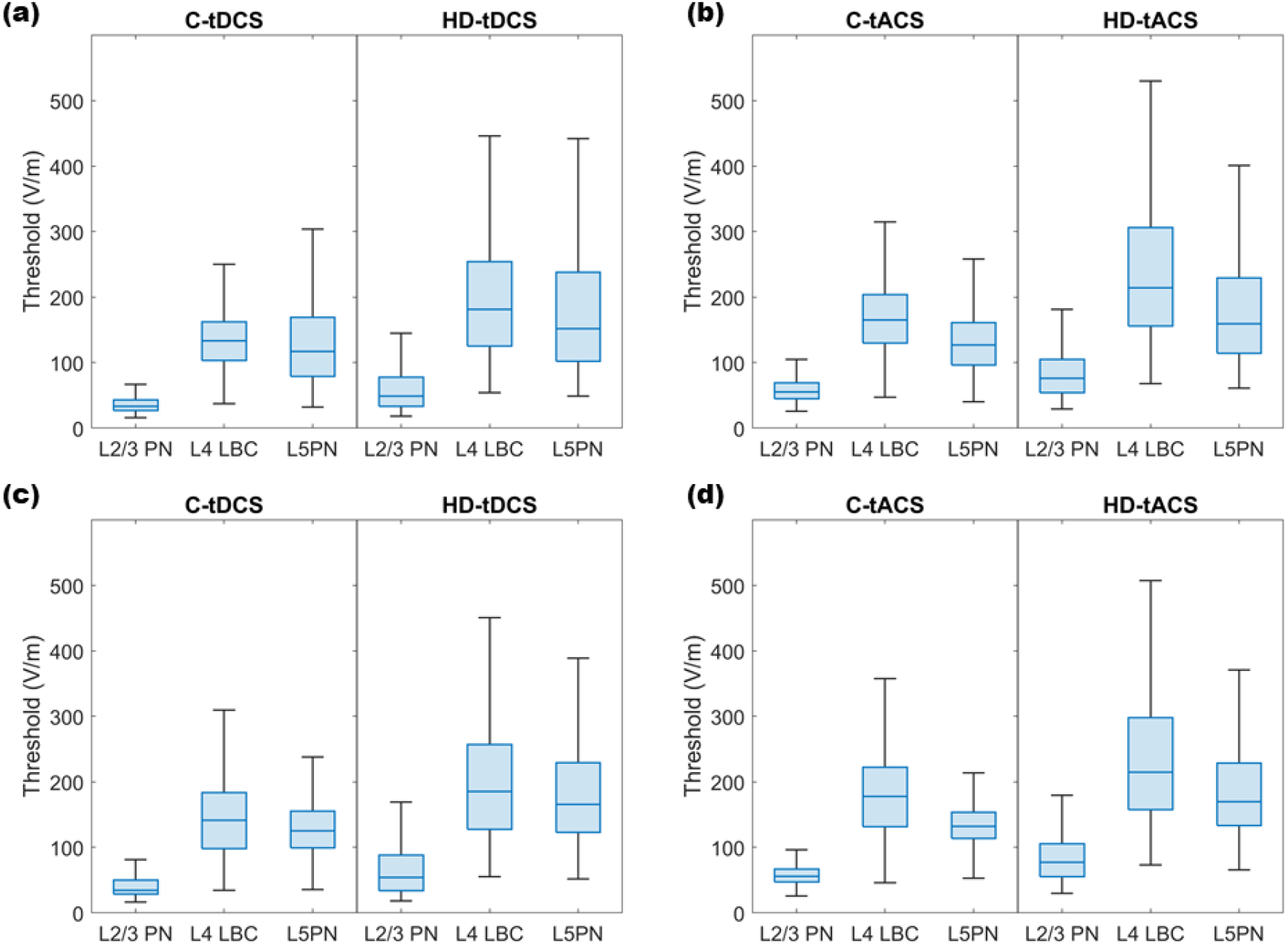
Thresholds for neurons populated in the ROI. The thresholds of L2/3PN, L4 LBC, and L5 PN in the unrotated case: (a) comparison between C-tDCS/HD-tDCS (b) comparison between C-tACS/HD-tACS. Further, in the rotated case: (c) comparison between C-tDCS/HD-tDCS (d) comparison between C-tACS/HD-tACS. Note that the rotated neurons’ threshold is the mean threshold of three independent rotation trials.

Moreover, in all C-/HD-tDCS/tACS and both the rotated/unrotated cases, a higher excitation threshold area was observed on the bank, which implies that such an area may be stimulated with a weaker EF. In addition, a lower threshold area was observed on the crown, implying that the area may be stimulated with a stronger EF (Figure 5–8). We observed this significant trend across all three types of neurons with all types of stimulations and in both rotated/unrotated cases, despite the differences in the cell localization depth. On the other hand, we found small differences among cell types. The lower threshold was primarily on top of the gyrus, but the specific locations of the lower and higher threshold areas differed somewhat. In addition, a higher threshold areas of L4 LBC and L5 PN were observed more broadly within the sulcal wall, as neurons’ morphologies and locations in the cortical layer differ. Moreover, a difference was observed between tDCS and tACS. In the unrotated case and postcentral gyrus, L3 with C/HD-tDCS showed a lower threshold area in the anterior bank and higher threshold area in the posterior bank (Figure 5). In contrast, L3 PN with C/HD-tACS showed a higher threshold area in both the anterior and posterior banks (Figure 6). In the rotated case, a lower threshold area in the pre/post-central gyrus of L2/3 PN with C-tDCS was much broader than that with C-tACS.

**Figure 5.**
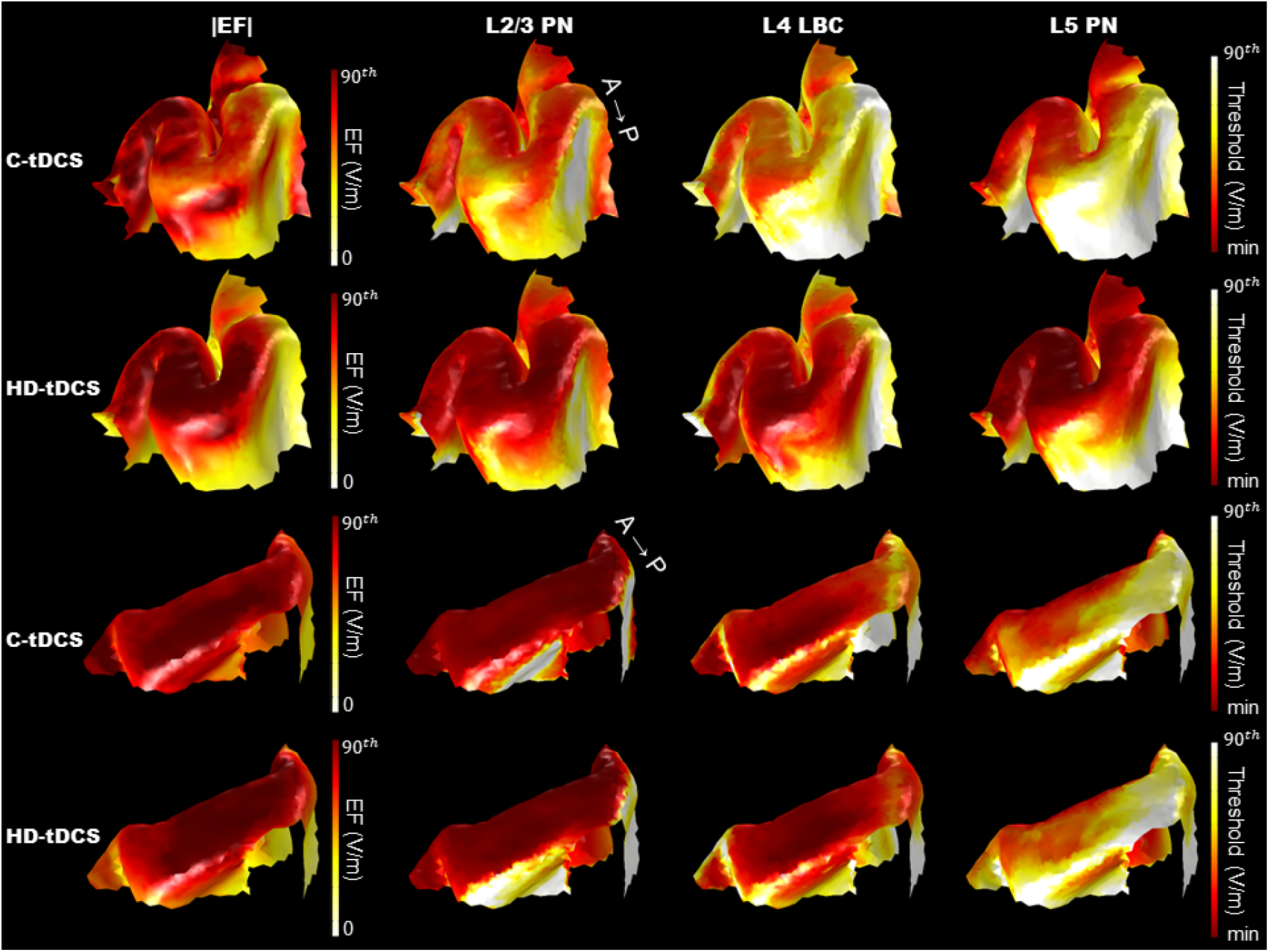
|EF| and threshold distribution of neurons stimulated with C-/HD-tDCS in unrotated case. The thresholds of neurons in the precentral gyrus are shown in the 1^st^ and 2^nd^ rows; those in the postcentral gyrus are depicted in the 3^rd^ and 4^th^ rows. The left column depicts the |EF| distribution from zero to the 90^th^ percentile. The remaining three columns depict L2/3 PN, L4 LBC, and L5 PN’s threshold distribution from zero to the 90^th^ percentile.

**Figure 6.**
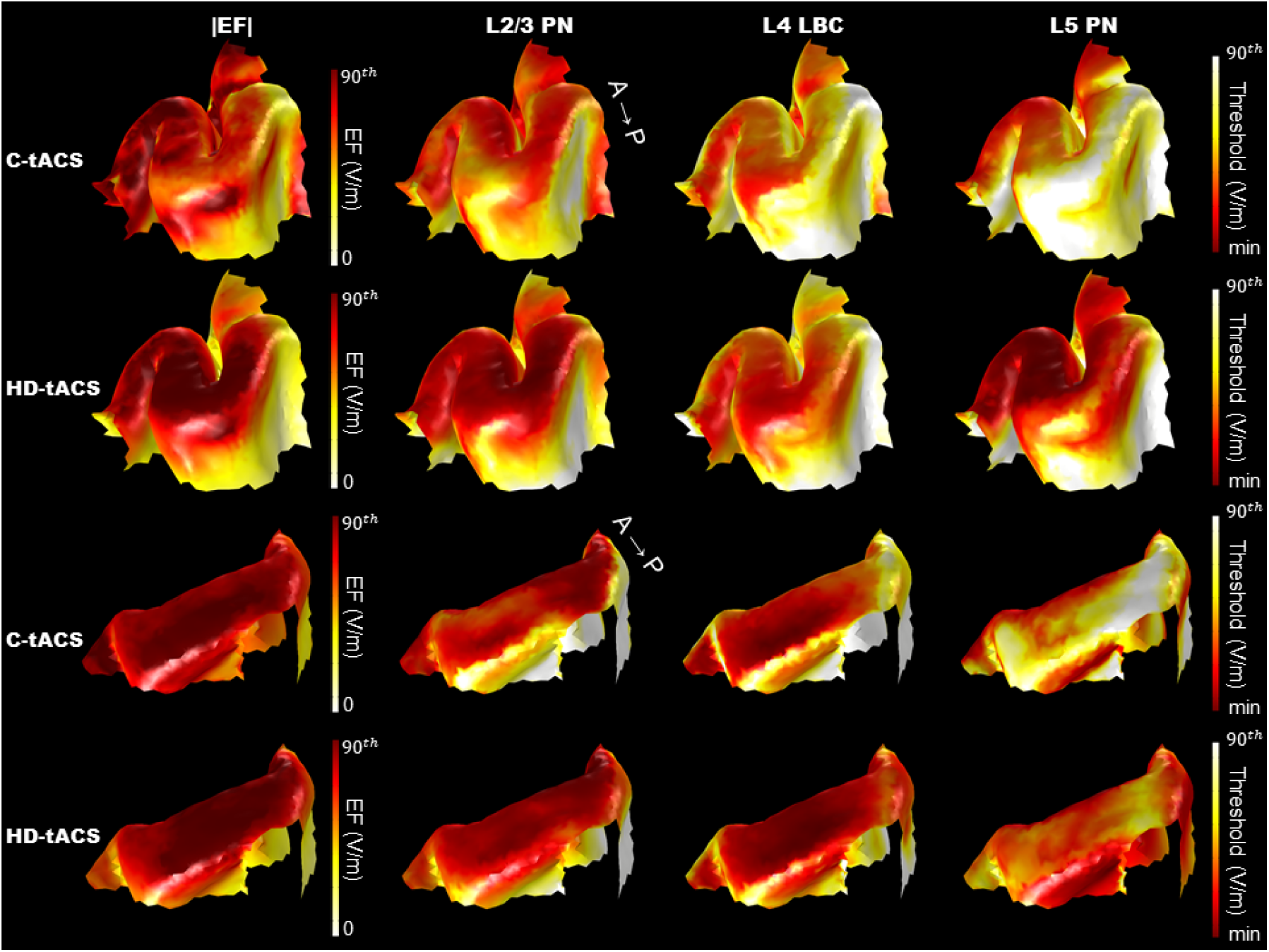
|EF| and threshold distribution of neurons stimulated with C-/HD-tACS in the unrotated case. The thresholds of neurons in the precentral gyrus are shown in the 1^st^/2^nd^ rows and those in the postcentral gyrus are depicted in the 3^rd^/4^th^ rows. The left column depicts |EF| distribution from zero to the 90^th^ percentile. The remaining three columns depict L2/3 PN, L4 LBC, and L5 PN’s threshold distribution from zero to the 90^th^ percentile.

When the rotated and unrotated cases were compared, we observed a notable pattern in the threshold distribution in the gyrus in the rotated case (Figure 7–8). In this case, even in adjacent regions, the threshold differed according to the rotation angles. This is because rotating neurons may change the location of the axon terminal and the |EF| applied to an axon may depend upon the latter’s location. Thus, a mottled pattern of threshold distribution was observed in the rotated, but not the unrotated case. Further, the mottled pattern was more notable in L5 PN than in L2/3 PN and L4 LBC. Because of the L5 PN’s greater depth, we believe the |EF| in layer 5 was weaker than in layers 1-4. The different |EF| applied to L5 PN and its morphology may have caused the distinct pattern.

**Figure 7.**
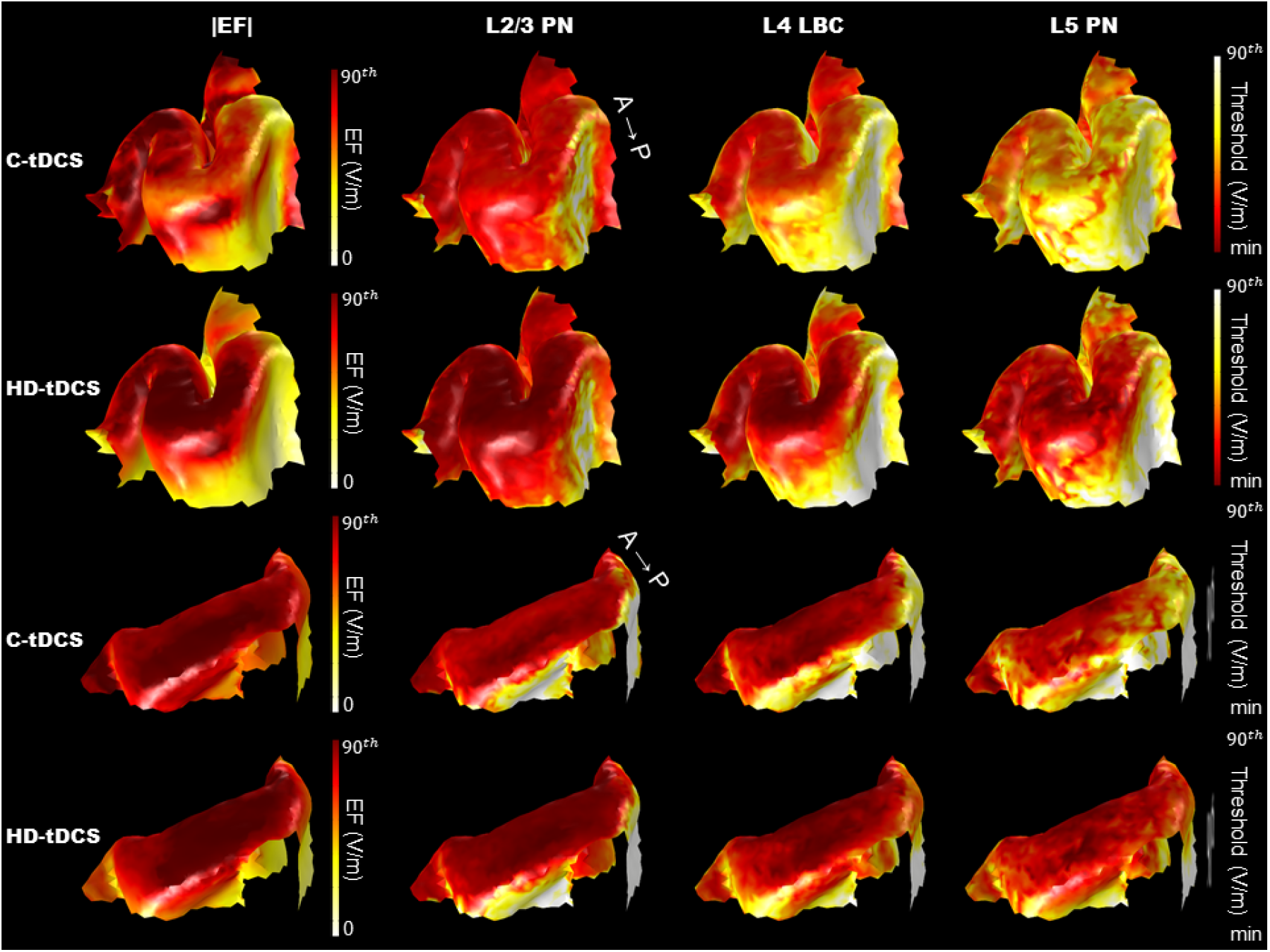
|EF| and threshold distribution of neurons stimulated with C-/HD-tDCS in the rotated case. The thresholds of neurons in the precentral gyrus are shown in the 1^st^/2^nd^ rows and those in the postcentral gyrus are depicted in the 3^rd^/4^th^ rows. The left column depicts |EF| distribution from zero to the 90^th^ percentile. The remaining three columns depict L2/3 PN, L4 LBC, and L5 PN’s threshold distribution from zero to the 90^th^ percentile.

### 3.3 Relation between |EF| component and threshold

Given the consistency in the |EF| and threshold results (Figure 5–8) and controversy over the relation between |EF| and threshold, we investigated the degree of correlation between |EF|/|RF| and the threshold with the Pearson correlation coefficient. It is known that higher |EF| generally induces a lower threshold and the converse, which indicates that the threshold is inversely proportional to |EF|. Further, we found a similar trend between |EF| and the threshold distribution in both the unrotated (Figures 5–6) and rotated case (Figures 7–8). Thus, we investigated the relation between the inverse of the threshold and |EF|/|RF|. Overall, the threshold was correlated more highly with |EF| than |RF| and the degree of correlation differed depending upon morphology and stimulation types. However, there was still a less significant difference in the correlation coefficient between the rotated and unrotated cases (Figure 9), and the correlation varied among the different neuron types. For example, the correlation coefficient of L5 PN was much lower than that of the other neurons. This may be because of the cortical layer depth, as layer 5 is much deeper in the cortex than layers 1-4 and thus, less of the EF reaches layer 5. Moreover, it may be attributable to the L5 PN neurons’ different morphology. As mottled patterns were observed notably in the rotated case of L5 PN, the L5 PN’s difference in PCC value between the rotated and unrotated cases was greater than the difference between the rotated and unrotated cases found in other types of neurons. In all C-/HD-tDCS/tACS, the PCC values of |EF|/|RF| in the rotated case were higher than those in the unrotated case. Moreover, in both the tDCS/tACS cases and unrotated/rotated cases, HD-tDCS/tACS demonstrated a higher correlation across the cell types, which may be attributable to the focality of HD-tDCS/tACS. Further, in both the rotated and unrotated cases, a difference in the PCC values of |EF|/|RF| was found in tDCS and tACS. The different PCC values according to neuron type, rotated or unrotated case, and stimulation montage show the various neuronal responses between them.

**Figure 8.**
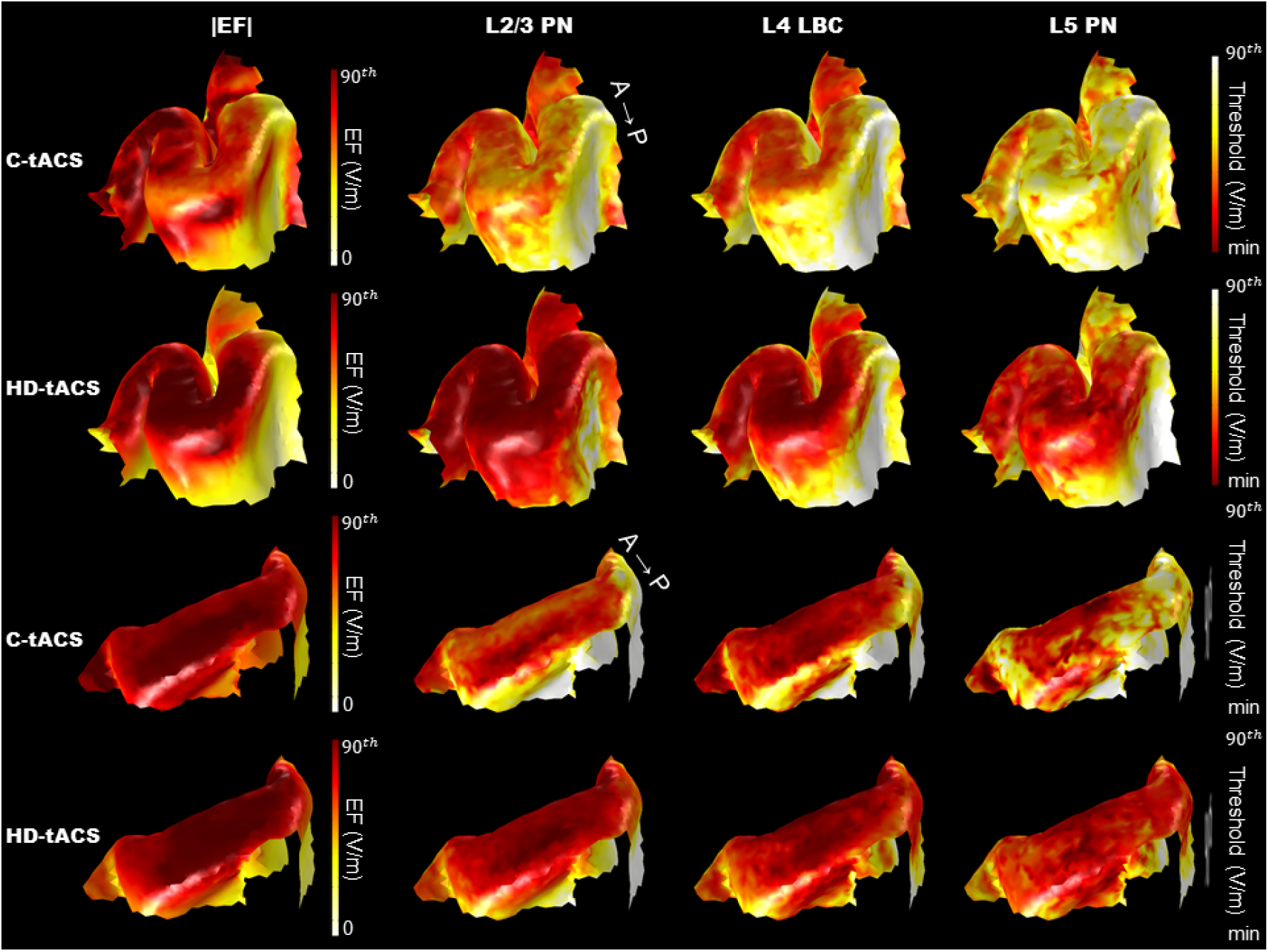
|EF| and threshold distribution of neurons stimulated with C-/HD-tACS in the rotated case. The thresholds of neurons in the precentral gyrus are shown in the 1^st^/2^nd^ rows and those in the postcentral gyrus are depicted in the 3^rd^/4^th^ rows. The left column depicts |EF| distribution from zero to the 90^th^ percentile. The remaining three columns depict L2/3 PN, L4 LBC, and L5 PN’s threshold distribution from zero to the 90^th^ percentile.

**Figure 9.**
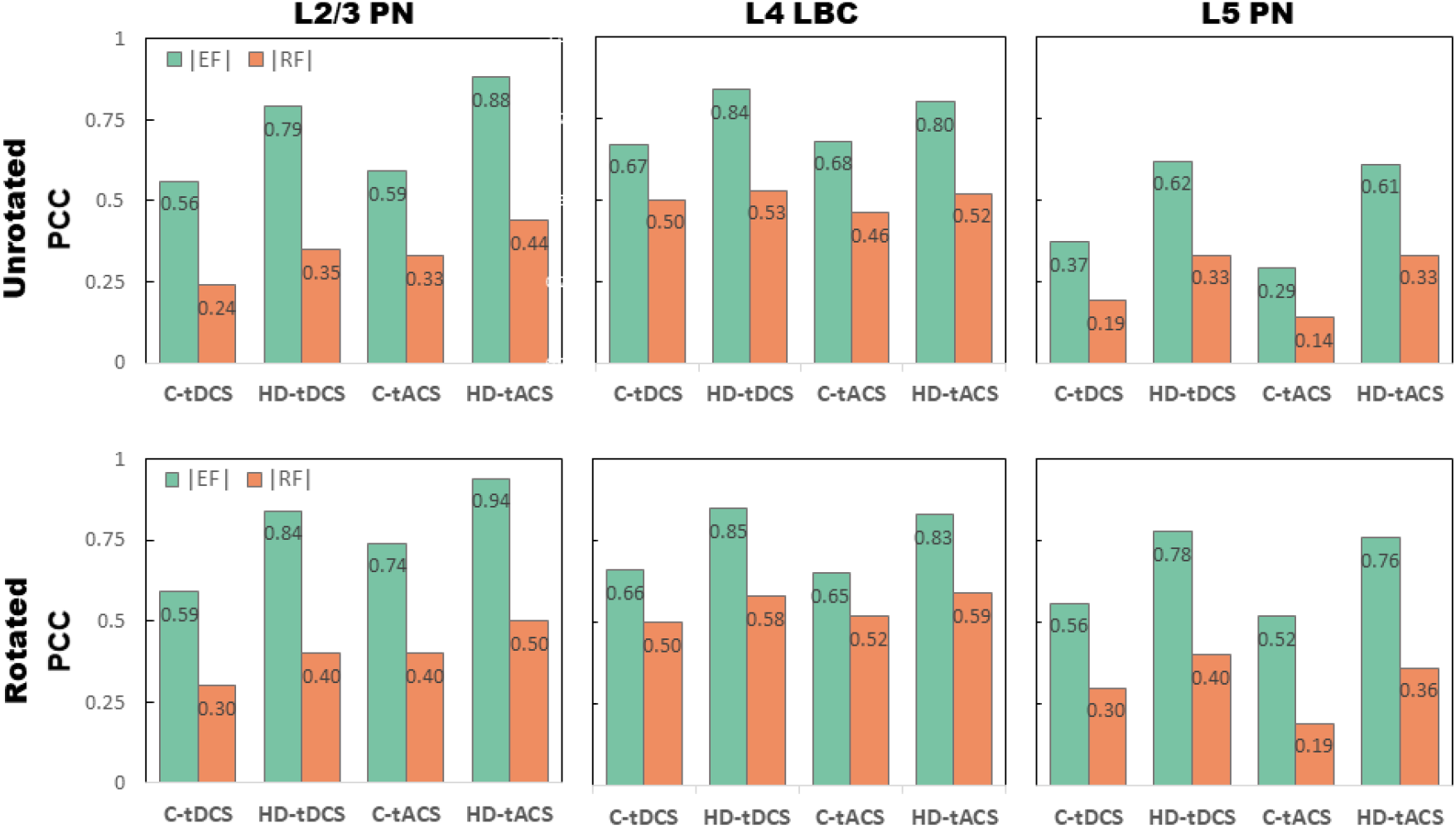
Correlation coefficient between |EF| and |RF|, and inverse of neurons’ thresholds. We estimated the correlations (PCC) between |EF| (green) and |RF| (orange) against the inverse of the threshold. The top row is the correlation in the unrotated case and the bottom row is in the rotated case.

## 4. Discussion

Various previous studies have investigated the determining factor of a stimulus-induced EF on neuronal excitability attributable to NIBS [16, 19, 28, 29, 31–37]. For TMS, Aberra et al. [29] constructed a multi-scale model using an arborized axon and found a higher correlation between neuronal activation and EF. In contrast, in a previous paper, our group reported a higher correlation between neuronal activation and RF when we applied tDCS [28]. Although this controversy may be attributable to different types of electric versus magnetic stimuli, we hypothesize that the different types of neuronal models, particularly arborized axons and idealized straight axons, may be the main factor in this controversy. Therefore, we constructed a multi-scale model using the arborized axon in Aberra’s study for tES and investigated the determining factors of cortical excitability tES causes. Here, we found that the excitation thresholds were correlated highly with |EF| compared to |RF|, which is consistent with Aberra et al.’s TMS report [29]. There are several differences between neuronal models with arborized and straight axons; in our previous report using straight axons, the neuronal morphologies were acquired from the cat visual cortex and then the neurons were scaled to the human cortex’s dimensions. Thereafter, we added idealized straight axons that stretched to the white matter. When we applied uniform EF, RF activated the neuronal models with straight axons but TF elicited little activation. In contrast, the neuronal model that Aberra et al. constructed had different morphologies acquired from the somatosensory cortex from rats that both RF and TF activated.

According to Aberra et al.’s findings [29], the degree of neurons’ sensitivity to |RF| depended upon the degree of an axon’s arborization. They found that the straight axon model had a greater preference for the downward orientation of EF, while the arborized axon model had a smaller preference for the EF’s orientation. Thus, it could be said that the straight axon model’s greater preference for |RF|, which is the strength of the downward EF, caused a higher correlation between the |RF| and threshold. It is accepted largely that EF flowing into a neuron parallel to a straight axon contributes primarily to a neuron’s polarization [28, 34, 35, 57]. To confirm the axon morphology’s importance, we replaced the arborized axons with a single straight axon without modifying the neuron models in any other way (Figure 10). The straight axon included myelination and a node and was aligned normal to the cortical surface within the head model. Such neuron models were populated in the ROI and then simulated. We found that neuronal models with straight axons increased the correlation between excitation thresholds and |RF| compared with arborized axons; thus, both EF and RF showed comparable degrees of correlation with the threshold. Unlike straight axons that have a single point of axon terminal, arborized axons, which extend in all directions, may allow neurons to respond to all directions of EF [29] because our neurons’ responses to EF were initiated in the axon terminal rather than the soma [7]. In addition, dendrites close to the positive field serve as an effective sink and those close to the negative field as an effective source [9]; thus, morphological differences may change the total current flow of the sink and source effects. Therefore, it is clear why the neuronal model with arborized axons we used in this work yielded a higher correlation with EF rather than RF.

**Figure 4.**
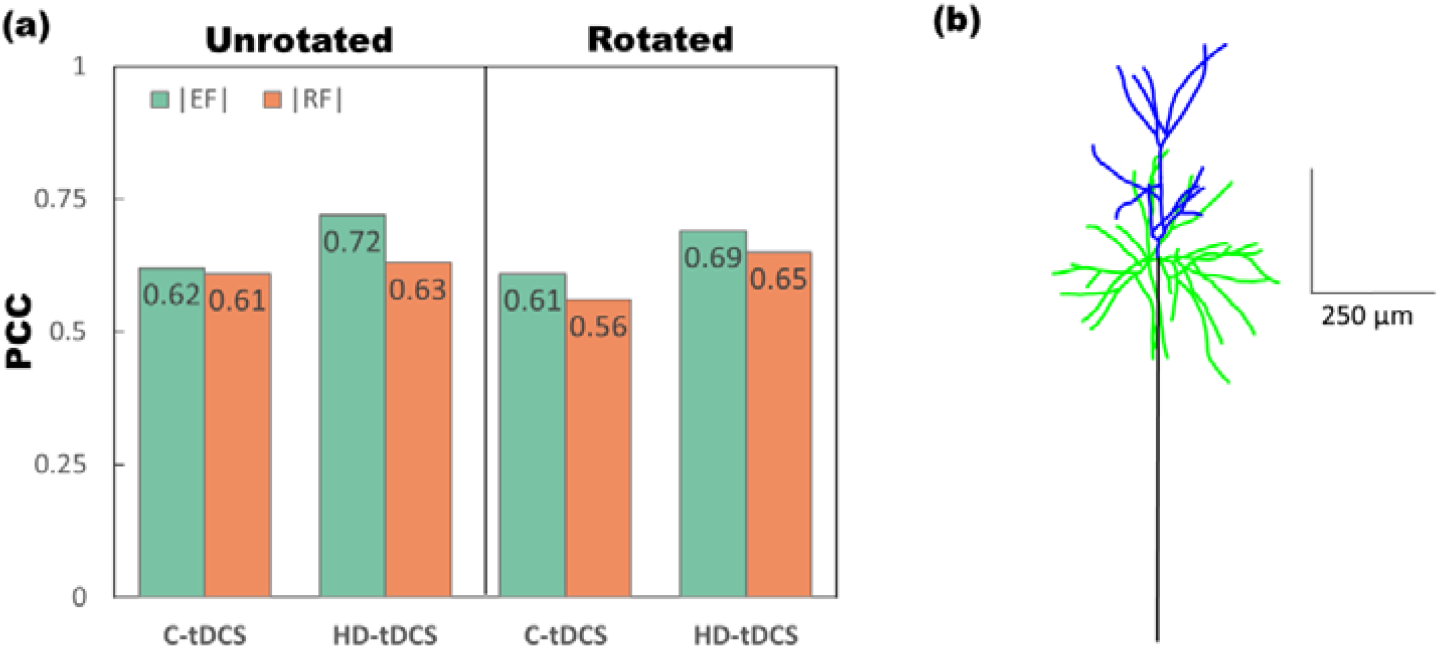
The correlation coefficient between |EF| and |RF|, and inverse of L2/3 PNs’ thresholds with a single straight axon, and morphology of the neuron. We replaced axon arbors with a single straight axon (b), and then estimated the correlations between |EF| (green) and |RF| (orange) against the inverse of the thresholds (a). The left box is the correlation in the unrotated case and the right box is in the rotated case.

Here, we stimulated neurons with two waveforms: a square wave (tDCS) and a sinusoidal wave (tACS). Because sinusoidal wave current fluctuates, tACS may apply a lower cumulative amount of current to neurons than tDCS. Thus, the excitation thresholds of neurons stimulated with tACS appear to be higher than those of neurons with tDCS. In addition, a difference between tDCS and tACS was found in the threshold distribution on the gyrus and the extent of correlations, although the difference was small. Such a small difference implies that the waveform of the current itself determines the total amount of current applied to the neurons, but cannot change the pattern of the neuronal response to electrical stimulation. In addition, a difference was found between both rotated and unrotated cases. The mottled pattern of the threshold distribution was observed in the rotated case only (Figures 7–8), which implies that different morphology may cause different threshold distributions. Further, the extent of the correlation differed according to the rotated and unrotated cases, and the difference was greater in L5 PN than in the other types of neurons. Overall, the extent of correlation between |EF|/|RF| varied according to the stimulation montage, waveform, and the rotated or unrotated case. However, the trend in which the neuron’s threshold was correlated more with |EF| than |RF| was consistent. This may be because there is no optimal direction of the EF in an azimuthal direction in our neuron model.

In addition to neurons’ axon morphology, their biophysical properties should be considered carefully. Another difference between our previous and current studies was the neurons’ biophysical properties; thus, such a difference in both morphology and biophysical properties may influence all of the results. The biophysical mechanisms vary according to the animal from which the model originated, or the experimental data on which the neuron model was based. The different biophysical mechanisms included in the neuron model may contribute to the different estimations of ion flux and intracellular current. Thereby, we may deduce that different morphological and biophysical properties may contribute to the conflicting results in previous reports and our work.

In our multi-scale model, we constructed three different types of neuronal models with different morphologies and properties. When we applied uniform EF, L4’s thresholds for RF and TF were higher than those of the other cell types, while the L2/3 PN had a larger threshold difference between RF and TF. These differences may be attributable to the morphological difference among cell types. L4 LBC has a smaller axon diameter than other cell types, which is likely to cause a higher threshold [5], and L2/3 PN has a morphologically asymmetric and elongated axon, while L4 LBC has spherically symmetric and highly arborized axons (Figure 2). We believe that these PNs’ morphological properties may explain L2/3 PN’s larger threshold difference between uniform RF and TF. Similarly, morphological features were also reflected in the threshold for realistic EF, and thereby, the correlations between EF and RF against the threshold. The threshold for realistic EF (Figure 5) was much higher than that for uniform EF, because the uniform EF applied was 1mV/mm, but realistic EF varied in a range from 0-0.4 mV/mm (C-tDCS/tACS) to 0-0.2mV/mm (HD-tDCS/tACS) (Figures 1c and 1d). Further, the complex geometry caused the direction of the EF applied to the neurons to vary. Because of these large differences, it is more desirable to use a multi-scale model to investigate cortical neurons’ responses to EF, which are the cellular targets for NIBS.

Although we introduced an advanced model for electrical stimulation in this work, several limitations of our models on the neuron and cortical scales should be recognized. With respect to the neuron scale, we adopted rat neurons obtained from the Blue Brain Project [46, 47] and modified them by adding myelination and scaling them to human neuron size [7]. We observed that the cortical neuronal response depended upon neuronal morphology, particularly axonal morphology. However, both this work and previous studies have adopted animal models and animal model neuron characteristics are highly likely to differ from human model’s characteristics. Thus, it is imperative to incorporate human neuron models. Another limitation is that our current model does not consider synaptic input from local and long-range intracortical connections. Electrical stimulation activates cortical neurons directly, and simultaneously, synaptic inputs themselves (recurrent connectivity) activate them and other connected neurons indirectly [58, 59]. Thus, in real neurons, excitatory inputs are expected to reduce the threshold and inhibitory inputs to increase it. To consider synaptic inputs, network models have been proposed to reproduce the indirect activation of neurons that synaptic inputs induce [60, 61]. However, multi-scale modeling studies have focused on the direct activation of neurons the EF induces [28, 29]. We determined the neuronal response to the EF and found that its magnitude itself is a crucial factor in cortical activity rather than its orientation. Further, we demonstrated that the determining factors in neuronal responses depend upon the neuron models’ morphological features. Although our investigation considers only the direct activation of cortical neurons, our findings should be taken into account synergistically in network models that consider both direct and indirect activation, which should be explored in future work.

With respect to the cortical level, we assumed isotropic conductivity within the gray and white matter, and calculated the EF in the head model. As a realistic head model obtained from MRI may provide an accurate estimation of the EF’s distribution, much research has investigated inter-subject variability attributable to individual differences in skull thickness, the white matter’s anisotropic conductivity, and the cortex’s anatomical geometry [20, 22, 62–66]. However, in this work, neurons were located within the gray matter and the white matter’s anisotropic conductivity does not affect the EF applied to neurons, although skull properties and anatomical geometry may affect the EF magnitude and orientation highly [22, 24, 43]. Given our observation of the EF magnitude’s contribution to neuronal responses, we should incorporate neuron models with various realistic head models, particularly those that reflect inter-subject variability in skull properties, the white matter’s anisotropic conductivity, and anatomical geometry.

To the best of our knowledge, most NIBS studies have investigated the motor cortex only because of the clear outcomes, such as MEP (motor-evoked potential). With such reasoning, our investigation was confined to an ROI in the hand knob area in the precentral gyrus (motor cortex) and its adjacent part of the postcentral gyrus. We note that exploration of the entire brain may be less tractable because of the greater computing resources required to simulate a myriad of neurons. Although our investigation focused on the motor cortical area, it is believed that our findings may be applied easily to other areas. Computational studies have a great advantage, in that other targeted areas with various montages can be applied cost-effectively, and thus, the EF distribution can be estimated [65, 67, 68]. Moreover, computational head modeling based upon MRI has advantages, in that it can predict the EF on the targeted area precisely and optimize the electrode stimulation montage [24]. Thus, our findings may help researchers determine optimal cellular targets and cortical areas that NIBS affects in future multi-scale modeling studies.

In conclusion, we observed that the EF magnitude was a stronger factor in cortical activity than that of RF, a normal component of EF in all electrode montages and waveforms, when EF was applied to neurons with axon arbors. Further, we demonstrated that this result may change when a single straight axon is introduced in place of an arborized axon. For neurons with straight axons, both RF, as well as the EF magnitude, were imperative factors. Taking into account previous reports [28, 29] and our findings, we conclude that the neuronal response depends somewhat upon the neuron models that are incorporated in the head model. Unfortunately, previous studies and our work have used models that are realistic, but are derived from animal neurons and scaled to the size of human neurons. Thus, precise electrophysiological and morphological properties of cortical neurons that are similar to actual human neurons are essential for precise estimation and better understanding of the NIBS mechanism.

## Acknowledgments

This work was supported by the Institute of Information & Communications Technology Planning & Evaluation (IITP) grant funded by the Korea Government (No, 2017-0-00451), and the ‘2021 Joint Research Project of Institutes of Science and Technology.’

